# The heme-responsive PrrH sRNA regulates *Pseudomonas aeruginosa* pyochelin gene expression

**DOI:** 10.1101/2023.01.19.524833

**Authors:** Tra-My Hoang, Weiliang Huang, Jonathan Gans, Evan Nowak, Mariette Barbier, Angela Wilks, Maureen A. Kane, Amanda G. Oglesby

## Abstract

*Pseudomonas aeruginosa* is an opportunistic pathogen that requires iron for growth and virulence, yet this nutrient is sequestered by the innate immune system during infection. When iron is limiting, *P. aeruginosa* expresses the PrrF1 and PrrF2 small regulatory RNAs (sRNAs), which post-transcriptionally repress expression of non-essential iron-containing proteins thus sparing this nutrient for more critical processes. The genes for the PrrF1 and PrrF2 sRNAs are arranged in tandem on the chromosome, allowing for the transcription of a longer heme-responsive sRNA, termed PrrH. While the functions of PrrF1 and PrrF2 have been studied extensively, the role of PrrH in *P. aeruginosa* physiology and virulence is not well understood. In this study, we performed transcriptomic and proteomic studies to identify the PrrH regulon. In shaking cultures, the pyochelin synthesis proteins were increased in two distinct *prrH* mutants compared to wild type, while the mRNAs for these proteins were not affected by *prrH* mutation. We identified complementarity between the PrrH sRNA and sequence upstream of the *pchE* mRNA, suggesting potential for PrrH to directly regulate expression of genes for pyochelin synthesis. We further showed that *pchE* mRNA levels were increased in the *prrH* mutants when grown in static but not shaking conditions. Moreover, we discovered controlling for the presence of light was critical for examining the impact of PrrH on *pchE* expression. As such, our study reports on the first likely target of the PrrH sRNA and highlights key environmental variables that will allow for future characterization of PrrH function.

**Importance:** In the human host, iron is predominantly in the form of heme, which *Pseudomonas aeruginosa* can acquire as an iron source during infection. We previously showed that the iron-responsive PrrF sRNAs are critical for mediating iron homeostasis during *P. aeruginosa* infection; however the function of the heme-responsive PrrH sRNA remains unclear. In this study, we identified genes for pyochelin siderophore biosynthesis, which mediate uptake of inorganic iron, as a novel target of PrrH regulation. This study therefore highlights a novel relationship between heme availability and siderophore biosynthesis in *P. aeruginosa*.

## Introduction

*P. aeruginosa* is a versatile environmental organism and opportunistic pathogen that can survive in a wide range of environments. As a pathogen, *P. aeruginosa* causes acute lung and blood infections in cancer patients and 10% of all hospital-acquired infections (1-4). *P. aeruginosa* also causes life-long chronic lung infections in individuals with cystic fibrosis (CF) and is a significant contributor to chronic wound infections in diabetics and surgical patients (5-7). To evade the immune system during infection, *P. aeruginosa* deploys numerous virulence factors, including exotoxin A (8-10), type three secretion (11-13), and redox-active phenazine metabolites (14-16).

*P. aeruginosa* can also form biofilms, or adherent communities encased in a self-produced exopolysaccharide (EPS) matrix, which protect the bacteria from immune assault during device-mediated (e.g. ventilator associated pneumonia) and chronic infections (17-19). *P. aeruginosa* is innately resistant to many therapeutic agents, and the emergence of multi-drug resistant (MDR) strains of *P. aeruginosa* leads to persistent infections, longer hospital stays, and increased mortality rates (20). Biofilm formation during chronic infection further complicates treatment due to increased tolerance of these communities against antimicrobials (21). Timely expression of virulence-related genes is essential for survival in the host, and *P. aeruginosa* regulates virulence-associated processes in response to a variety of environmental cues, including nutrient availability and quorum sensing factors (22). Understanding the regulatory pathways that mediate virulence trait expression may therefore reveal novel strategies for therapeutic intervention.

As with many other pathogens, *P. aeruginosa* requires metallonutrients for growth and virulence. *P. aeruginosa* has a particularly high requirement for iron, which plays a central role in metabolism, oxygen and redox sensing, protection from oxidative stress, and nucleic acid synthesis (23). To limit pathogen growth, the host restricts iron and other essential metals in a strategy called “nutritional immunity” (24). *P. aeruginosa* overcomes nutritional immunity through a variety of mechanisms, including the synthesis and uptake of two siderophores – pyoverdine and pyochelin – which scavenge the oxidized (ferric) form of iron (Fe^3+^) from the host iron sequestration proteins lactoferrin and transferrin (25-27). In reducing environments, *P. aeruginosa* acquires the reduced (ferrous) form of iron (Fe^2+^) via the Feo system (28). *P. aeruginosa* can also acquire iron from heme, representing the most predominant source of iron in the human body (29). Heme acquisition is mediated by the heme assimilation (Has) and *Pseudomonas* heme uptake (Phu) systems, which transport heme into the cytosol (30), and a cytosolic heme oxygenase HemO that cleaves the heme tetrapyrrole to yield biliverdin and inorganic iron for use in cellular processes (31). Several studies have suggested that siderophore-mediated iron uptake is critical for acute infections (26, 32), while ferrous and heme uptake become more prominent in chronic, biofilm mediated infections that are characterized by biofilm communities, steep oxygen gradients, and persistent inflammation (33-37).

Despite its essentiality, iron catalyzes the formation of reactive oxygen species via Fenton chemistry, leading to damage of membranes, proteins, and DNA. In *P. aeruginosa* and many other bacteria, the Ferric uptake regulator (Fur) when bound to cytosolic Fe^2+^ becomes an active transcriptional repressor of genes involved in iron uptake (38, 39). *P. aeruginosa* Fur also represses expression of two non-coding small RNAs (sRNAs) called PrrF1 and PrrF2 (40). The PrrF sRNAs function by complementary base-pairing with, and destabilization of, mRNAs coding for non-essential iron-containing proteins, resulting in what has been termed the “iron sparing response” (41, 42). Owing to the central role of iron in *P. aeruginosa* physiology, deletion of both the *prrF1* and *prrF2* genes results in a significant growth defect in low iron media, decreased production of quorum sensing molecules, increased susceptibility to tobramycin during biofilm growth, and attenuated virulence in an acute murine lung infection model (43-48).

The *prrF1* and *prrF2* genes are located in tandem on the *P. aeruginosa* chromosome, allowing for the transcription of a distinct third sRNA called PrrH (49). PrrH shares a promoter and transcriptional start site with PrrF1, yet its expression is also dependent on read-through of the *prrF1* Rho-independent terminator, *prrF1-prrF2* intergenic region, and *prrF2* sequence (**Fig. 1A**).

**Fig 1.**
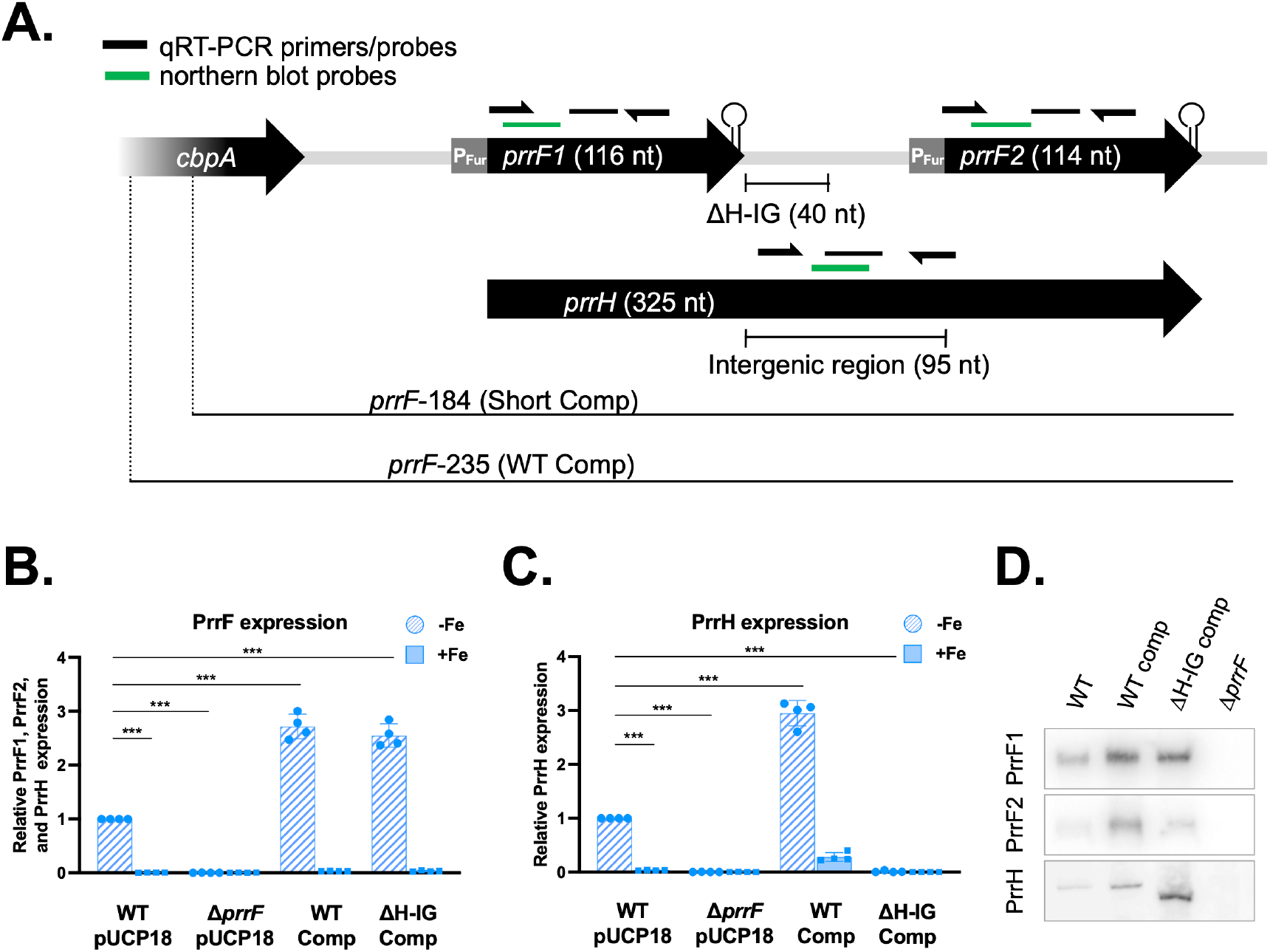
Characterization of transcripts produced by the Δ*prrH-IG* mutant. Organization of the *prrF* locus and design of complement plasmids (A). The WT comp includes the entire *prrF* locus plus 235 bp upstream of the *prrF1* promoter to ensure regulation of locus is uninhibited. The ΔH-IG comp also includes the 235 bp region upstream of the *prrF1* promoter but has 40 bp of the *prrF1-prrF2* intergenic region removed. The short comp contains the entire *prrF* locus but only includes 184 bp upstream of the *prrF1* promoter. Location of qRT-PCR primers and probes are indicated in pink (PrrH) and red (PrrF). The PrrF primers and probes cannot distinguish between PrrF1 and PrrF2. Northern blot probes are labeled in blue. For qRT-PCR (B,C), PrrF and PrrH transcription are shown as an average of 3 biological replicates, relative to WT PAO1 in low iron. Northern blot (D) is a representative from multiple experiments using radiolabeled DNA probes specific to each transcript.

Due to sharing a promoter with *prrF1*, transcription of PrrH is similarly repressed by iron (49). However, PrrH has also been shown to be regulated by heme (49, 50). Moreover, the PrrH sRNA contains a unique sequence, derived from the *prrF1-prrF2* intergenic region (**PrrH-IG, Fig. 1A**), that may be able to interact with and alter stability or translation of a distinct regulon of mRNAs. The entire *prrH* sequence, including the PrrH-IG region, is broadly conserved in *P. aeruginosa* clinical isolates, and the PrrH transcript is detected in clinical sputum samples, suggesting importance of this sRNA during infection (34). An inherent challenge of studying PrrH is separating its functions from those of the PrrF sRNAs, since it is not possible to transcribe PrrH without the *prrF1* and *prrF2* genes (**Fig. 1A**). Given the model that the PrrH-IG region is required for PrrH function, we previously generated a *prrF* locus allele with a deletion of the PrrH-IG region (Δ*prrH-IG*) to distinguish PrrH and PrrF functions. We found that the PrrH-IG sequence is not responsible for any of the previously identified phenotypes of the Δ*prrF* mutant (44). Thus, the role of PrrH in mediating iron and heme homeostasis to date remains unclear.

In the current study, we continued our analysis of the Δ*prrH-IG* mutant, as well as a distinct *prrH* mutant, to further investigate PrrH function. These strains were characterized by multiple approaches, including simultaneous RNAseq and proteomics analyses. Our results revealed genes for pyochelin biosynthesis as possible targets of PrrH regulation. We subsequently showed that *prrH* mutation led to increased expression of *pchE*, and we identified static growth conditions as more permissive of this regulation. We further found that controlling for light, which can be sensed by *P. aeruginosa* through the photoreceptor BphP, led to more consistent and robust repression of *pchE* by PrrH, suggesting this signal may have confounded previous PrrH regulation studies. Lastly, we show that heme represses expression of the *pchE* gene, though the precise role of PrrH in this regulation remains unclear. Overall, our data indicate that heme and PrrH affect expression of pyochelin siderophore biosynthesis, either by distinct or overlapping pathways.

## Results

### Characterization of *prrH* mutants

To investigate the functions of PrrH, we used the WT and ΔH-IG complementation system previously developed by our laboratory (**Fig. 1A**) (44). In this system, the Δ*prrF* mutant is complemented with either the entire *prrF* locus (WT-comp) or the *prrF* locus lacking 40 bp of the *prrF1*-*prrF2* intergenic region (ΔH-IG-comp) (**Fig. 1A**) *in trans* using the pUCP18 vector. Strains labeled as wild type (WT) and Δ*prrF* are the indicated PAO1 strains carrying the empty pUCP18 vector. As previously observed (44), quantitative real-time PCR (qRT-PCR) using the primers in **Figure 1A** shows that WT-comp expresses PrrF and PrrH in low but not in high iron M9 minimal medium (**Fig. 1B**). Furthermore, the PrrF, but not the PrrH, transcript is detected in the ΔH-IG comp strain, indicating this strain is a *prrH* mutant that can still express PrrF (**Fig. 1C**). Both PrrF and PrrH are expressed at about 3-fold higher levels in the complemented strains compared to the WT vector control, likely due to ectopic expression from the pUCP18 plasmid.

To confirm the qRT-PCR results, northern blot analysis was performed with probes specific for PrrF1, PrrF2, and PrrH as indicated in **Figure 1A**. As expected, the PrrF1 and PrrF2 transcripts were detected in RNA isolated from the WT-comp and the ΔH-IG-comp strains grown in iron-depleted medium (**Fig. 1D**). In agreement with the qPCR results, both complemented strains expressed higher levels of PrrF than the WT vector control (**Fig. 1D**). Because the PrrH probe anneals to the intergenic region adjacent to the deleted 40 bp (**Fig. 1A**), we were also able to determine that the ΔH-IG comp strain expresses a shorter PrrH transcript compared to that expressed in the WT vector control and WT-comp strain, indicating that the ΔH-IG allele produces a truncated PrrH sRNA (**Fig. 1D**).

We next characterized the transcripts of a distinct *prrH* mutant. This mutant was originally constructed as a *prrF* complementation plasmid that contained less sequence upstream of the *prrF1*/*prrH* transcriptional start site than the “WT comp” used in the above discussed studies. This plasmid, which we refer to as the “short-comp”, was designed to contain the entire *prrF1*-*prrF2* locus with only 184 bp upstream of the *prrF1/prrH* transcriptional start site, as compared to 235 bp upstream that is included in the WT-comp (**Fig 1A**) (44). Confirming our previous observation (44), qRT-PCR analysis shows that the PrrF, but not the PrrH, transcript is detected in RNA isolated from the short-comp strain (**Fig 2A, B**). We next performed northern blot analyses with probes specific for PrrF1, PrrF2, and PrrH. The short-comp strain exhibited lower levels of the PrrF2 sRNA and higher levels of the PrrF1 sRNA as compared to WT-comp (**Fig. 2C**). Additionally, a PrrH transcript was detected from the short-comp strain, but it was slightly shorter than the PrrH transcript produced by the WT strain (**Fig. 2C**). These observations were unexpected, as the short complement was designed to only lack sequence well upstream of the *prrF1/prrH* transcriptional start site. To investigate this further, we sequenced the short complement plasmid and discovered a rearrangement in the *prrF2* region, as shown in the cartoon in **Figure 2D**. Specifically, 27 nucleotides flanking the *prrF2* start site (from -12 to +25) were translocated to downstream of the *prrF2* Rho-independent terminator (**Fig. 2D**). This would, therefore, account for both the fainter PrrF2 transcript and truncated PrrH transcripts that were detected in the northern blots. While we were still able to detect a PrrH transcript in the short comp, the re-arrangement led to a PrrH transcript that was truncated in a distinct manner from the ΔH-IG mutant. Thus, we used both the short-comp and ΔH-IG *prrH* mutants to investigate PrrH regulation in this study.

**Fig 2.**
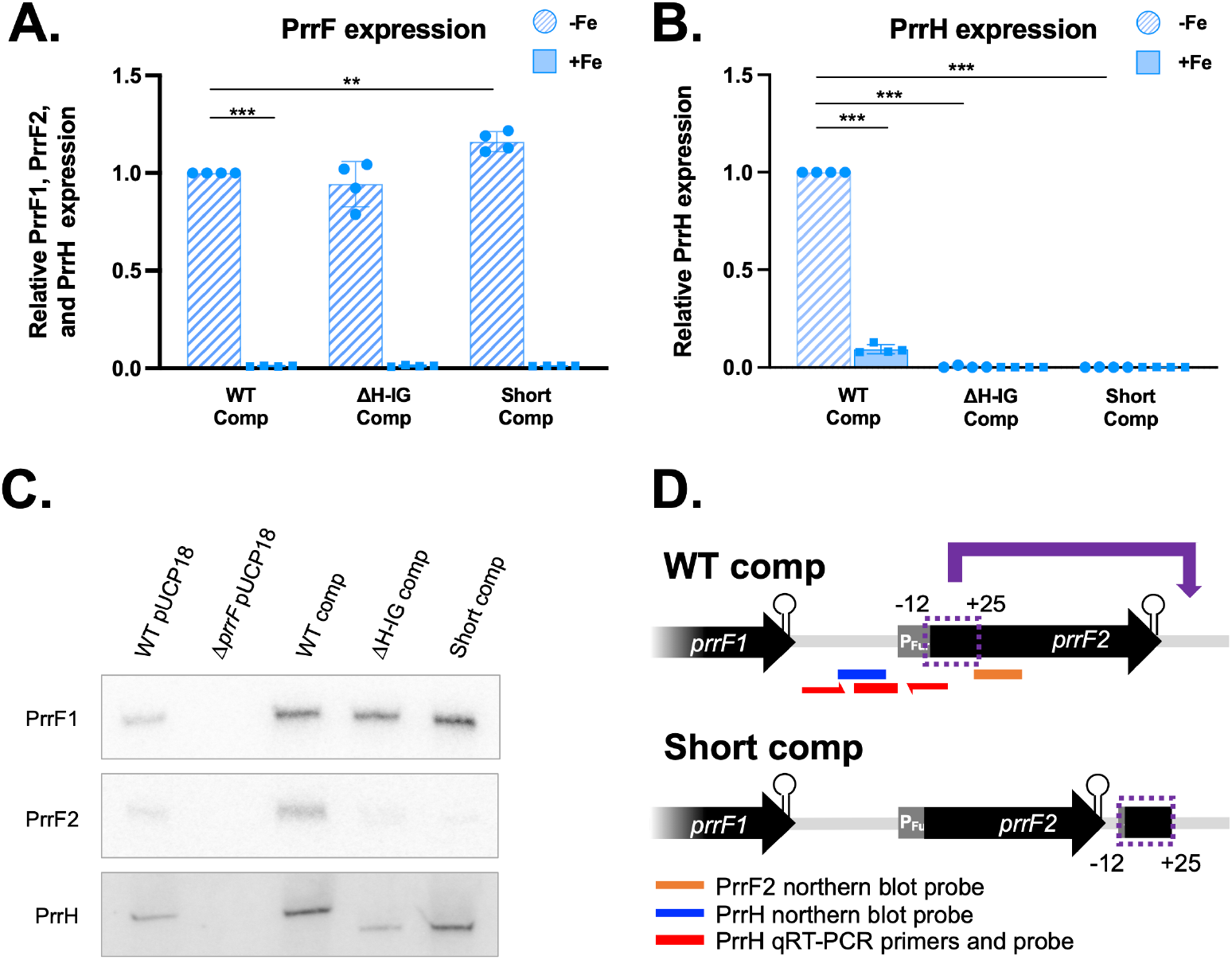
The short *prrF* complement has a rearrangement in *prrF2*. qRT-PCR analyses of PrrF (A) and PrrH (B) transcription are shown as an average of 3 biological replicates, relative to WT PAO1 in low iron. Northern blot (C) is a representative from multiple experiments using radiolabeled DNA probes specific to each transcript (PrrF1, PrrF2, and PrrH). RNAs were isolated from samples collected after 8 hours of aerobic growth in M9 media supplemented with 50 nM FeCl_3_ (−Fe, low iron) or 100 µM FeCl_3_ (+Fe, high iron) at 37°C. Panel D illustrates the re-arrangement that occurred in the short comp. Location of the northern blot probes used in 3C are shown in blue (PrrH) and orange (PrrF2). The qRT-PCR primers/probe for PrrH are shown in red. Sequencing was performed by Eurofins Genomics. Sequencing results were aligned and analyzed using MacVector software.

### Transcriptomic and proteomic analysis of PrrF and PrrH regulation in PAO1 reveals pyochelin biosynthesis as a potential PrrH target

Subsequent to validating the ΔH-IG-comp strain as a truncated *prrH* mutant, we performed RNAseq (51, 52) and label-free proteomics (53) on cultures of the WT vector control, Δ*prrF* vector control, WT-comp, and ΔH-IG-comp strains grown in M9 minimal media, with or without FeCl_3_ or 5µM heme supplementation. Five biological replicates for each group were processed and the resulting data analyzed as described in the Materials and Methods to generate a log fold change (LFC) for each RNA or protein in response to: 1) iron or heme supplementation of each strain, 2) deletion of the *prrF* locus by comparing the WT and Δ*prrF* vector controls, and 3) deletion of the H-IG sequence by comparing the WT-comp and ΔH-IG-comp strains. Full datasets are provided in the supplementary materials as **Datasets S1** (RNASeq) and **S2** (Proteomics). As expected, proteins involved in pyoverdine, pyochelin, and heme uptake, as well as their corresponding mRNAs, were repressed by iron in each of the strains, though the intensity of iron regulation varied somewhat amongst the individual strains, particularly at the protein level (**Supplementary Materials, Fig. S1A**). Also as expected, most of the known PrrF target mRNAs, and the proteins they encode, were activated by iron in the WT vector control, WT-comp, and ΔH-IG strain, and this regulation was reduced or eliminated in the *ΔprrF* vector control (**Supplementary Materials, Fig. S1A**), indicating that the plasmid-derived PrrF sRNAs function similarly to those transcribed from the chromosome. Regulatory patterns for these know iron-regulated genes and proteins varied slightly between the WT-comp and WT vector control, but they were comparable when comparing the WT-comp and ΔH-IG-comp strains (**Supplementary Materials, Fig. S1A**).

We next determined how loss of the H-IG sequence affected PAO1 gene expression by comparing the transcriptomes and proteomes of the ΔH-IG-comp strain, grown in low iron, to that of the WT-comp strain, also grown in low iron. This analysis revealed no statistically significant differences in the transcriptomes of these strains (**Supplementary Materials, Fig. S1B**), while the proteome of the ΔH-IG comp was substantially altered compared to that of the WT comp (**Supplementary Materials, Fig. S1C**). To validate observed changes in the PrrH-affected proteome, we conducted a subsequent proteomics experiment with the WT-comp, ΔH-IG-comp, and short-comp strains grown in M9 minimal medium with and without iron supplementation. As observed in the first experiment, proteins involved in pyoverdine, pyochelin, and heme uptake were similarly repressed by iron in all three strains, and PrrF-repressed targets showed similar effects in response to iron supplementation across all three strains (**Fig. 3A**), demonstrating that both *prrH* mutants exhibited iron and PrrF regulation similar to the WT strain. Also as observed for the first experiment, the proteomes of the ΔH-IG comp (**Fig. 3B**) and the short-comp (**Fig. 3C**) were significantly altered when compared to the WT-comp strain. Curiously, we also noted substantial differences in the proteomes of the short-comp and ΔH-IG strains (**Fig. 3D**). Moreover, proteins that were either upregulated or downregulated in each *prrH* mutant compared to the WT-comp in this dataset showed limited overlap with one another (**Fig 3E**, ΔH-IG-affected proteomes are represented by green circles, short-comp-affected proteomes are represented by yellow circles). Likewise, proteins that were either upregulated or downregulated upon H-IG deletion in the first (**Fig. 3F**, represented by pink circles) and second (**Fig. 3F**, represented by purple circles) proteomics experiments showed little overlap.

**Fig 3.**
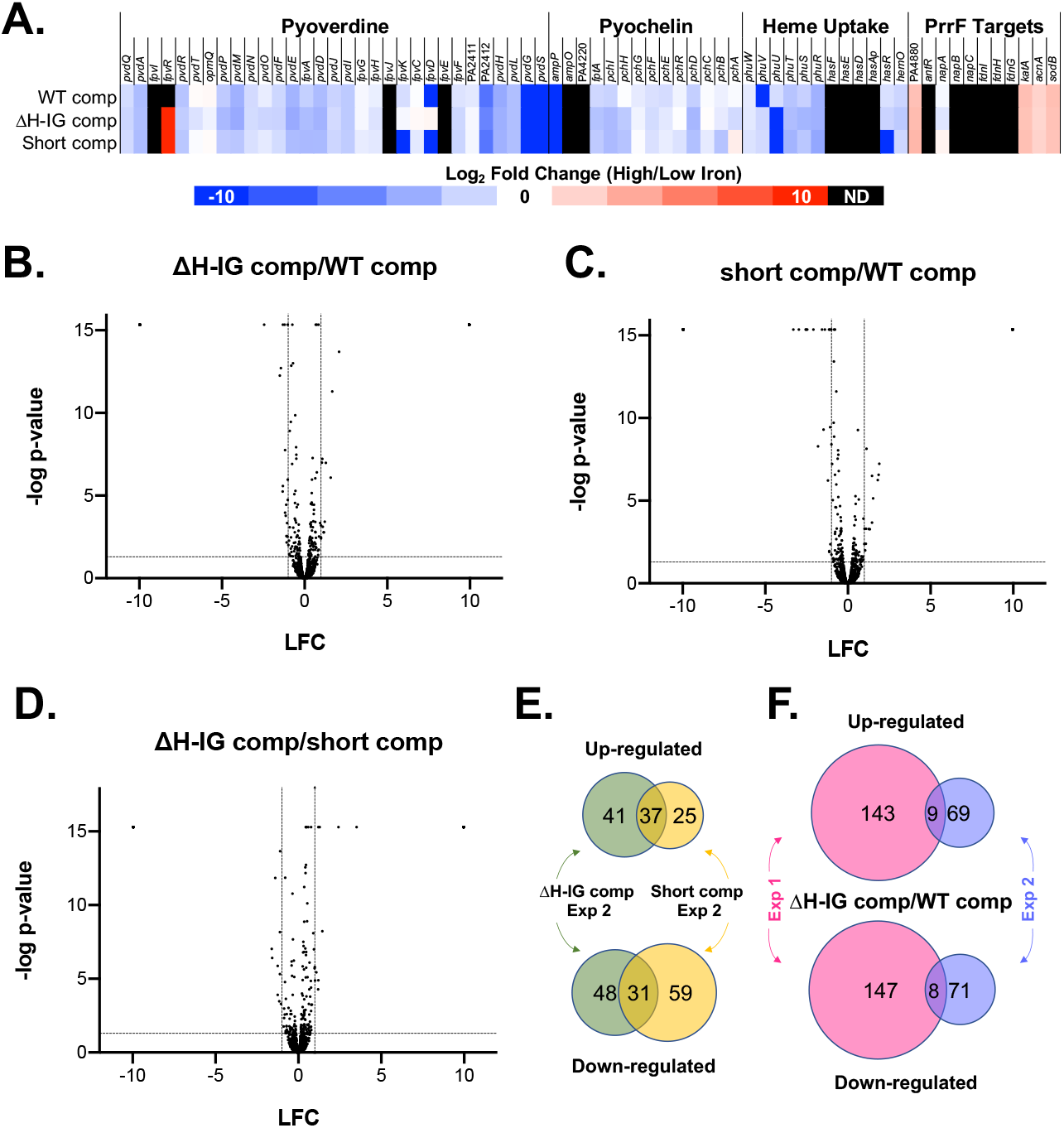
Short complement and ΔH-IG complement substantially alter the proteome. Proteomics results when comparing protein samples collected after 8 hours of aerobic growth in M9 media supplemented with 50 nM FeCl_3_ at 37°C. (A) Heatmap of a select group of iron-, heme-, and PrrF-regulated proteins shown as the log_2_ fold change of the abundance ratio between high iron and low iron. Undetected proteins are colored in black (ND). (B-D) Volcano plots comparing the protein abundance of the ΔH-IG comp versus WT comp (B), short comp versus WT comp (C), and ΔH-IG comp versus short comp (D). The log_2_ fold change is shown on the x-axes and the -log of the FDR p-value is on the y-axes. Horizontal dashed lines indicate FDR p=0.05 and vertical dashed lines indicate LFC = ±1. (E-F) Comparison of the dysregulated proteins in the ΔH-IG comp from experiments 1 and 2 are shown as Venn diagrams. (E) Panel E shows the overlap of the dysregulated proteins in the ΔH-IG comp and short comp, each compared to WT comp, from experiment (F) Panel F shows the overlap of the dysregulated proteins in the ΔH-IG comp compared to WT between experiments 1 and 2. To be included in the Venn diagram analysis, changes in protein levels must have demonstrated an FDR p-value <0.05 and -0.5≤ LFC ≥0.5.

Since the above datasets showed little consistency in robust PrrH effects, we sought to identify smaller yet more consistent regulatory effects of *prrH* mutation. To investigate this, we performed STRING network analysis on the proteins that were differentially regulated in any one of the three comparisons (ΔH-IG comp/WT comp from experiment 1, ΔH-IG comp/WT comp from experiment 2, and short comp/WT comp from experiment 2). To capture proteins that were weakly but statistically significantly affected in each experiment, we lowered the LFC cutoff value for proteins to be analyzed to 0.5 (−0.5≤ LFC ≤0.5). STRING network analysis revealed numerous dysregulated functions and pathways amongst the proteins affected in all three experimental comparisons (**Supplementary Materials, Fig. S2-S3**). However, many of these clusters were not consistently dysregulated. For example, the phenazine biosynthesis cluster was upregulated upon *prrH* mutation in the first experiment yet downregulated by *prrH* mutation in the second experiment. We also observed inconsistencies between comparisons when we considered how iron, heme, and PrrF affected genes in each of these clusters. For example, proteins in cluster A (sulfur metabolism) showed variable regulation by PrrH and were repressed in the *prrF* mutant but induced by iron (**Supplementary Materials, Fig. S2-S3**).

Despite the variations in the effects of PrrH amongst the different experiments, we identified two clusters in which the effects of PrrH on proteins involved in specific cellular functions were consistent across all three comparisons. Cluster M included proteins for 2-alkyl-4(1*H*)-quinolone biosynthesis which were reduced upon mutation of PrrH in all three comparisons (**Fig. 4**). In contrast, Cluster N was comprised of pyochelin proteins that were largely induced by PrrH mutations in all three comparisons (**Fig. 4**). Consistent with the analysis shown in **Figure S1**, none of the RNAs encoding these proteins were significantly affected by PrrH mutation in the RNASeq analysis of the first experiment (**Fig. 4**). Further analysis of Cluster M proteins showed that these proteins were also affected by *prrF* deletion, which is consistent with previous studies from our group (34, 48). Moreover, while iron repressed the levels of RNAs for the Cluster M proteins, heme did not affect levels of the Cluster M proteins (**Fig. 4**), suggesting they are not specifically regulated by PrrH. In contrast, proteomics and RNAseq showed no effect of PrrF on proteins in Cluster N (**Fig. 4**). Moreover, heme repressed proteins in Cluster N, but had no effect on the corresponding RNAs (**Fig. 4**), consistent with the lack of transcriptome effects observed upon mutation of the PrrH sRNA (**Supplementary Materials, Figure S1**). Therefore, we focused on the pyochelin biosynthesis genes and proteins in Cluster N as potential novel targets of the PrrH sRNA.

**Fig 4.**
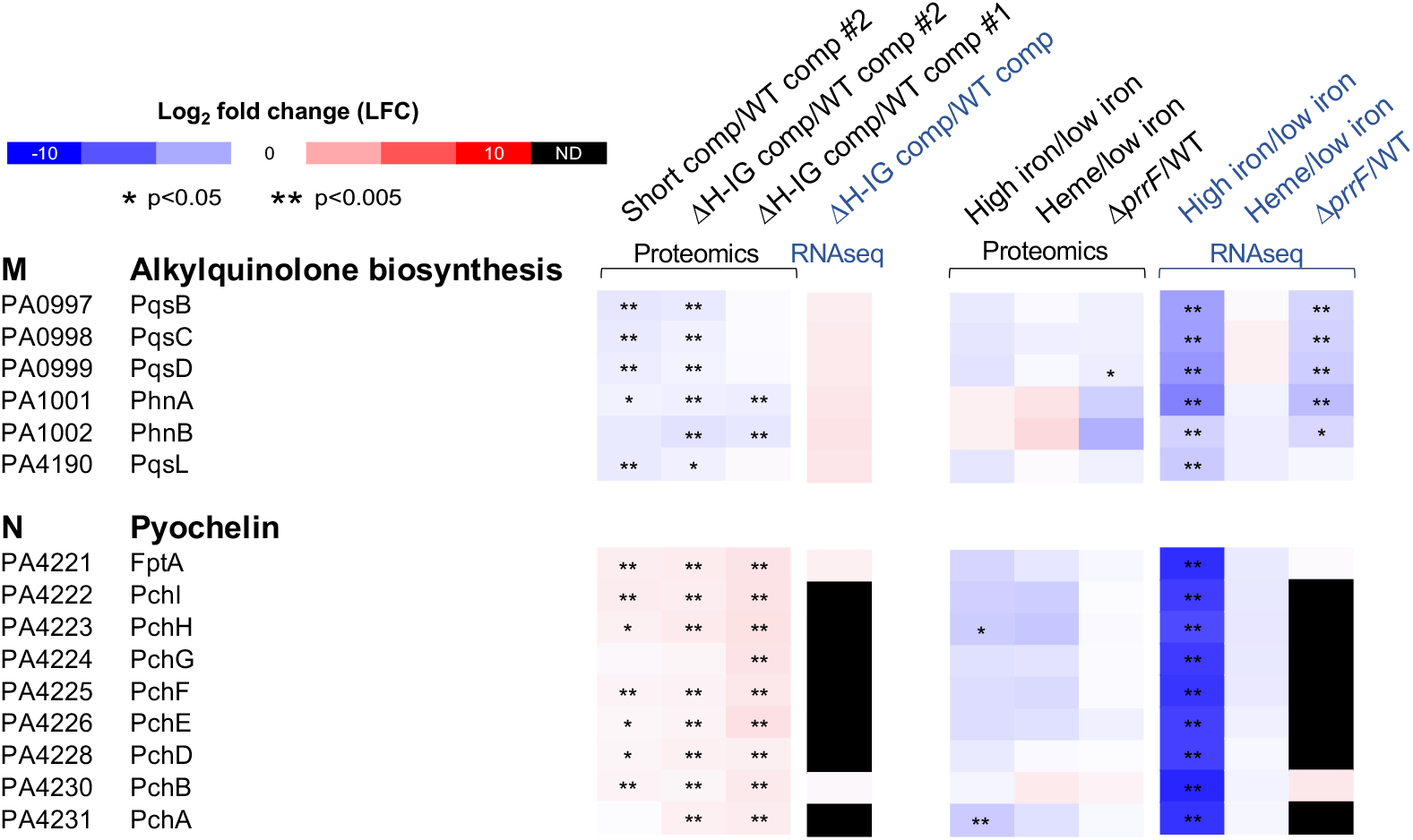
Proteins involved in alkylquinolone biosynthesis and pyochelin biosynthesis are dysregulated in *prrH* mutants. Expression data from proteomics and RNAseq are presented as heat maps where up-regulated proteins are indicated in red and those down-regulated are in blue. Undetected proteins are colored in black (ND). ** indicate p<0.005 and * indicate p<0.05.

### Static growth reveals potential for PrrH-mediated repression of the pyochelin biosynthesis *pchE* mRNA

sRNA-mediated repression of gene expression is most often mediated by pairing at or near the Shine Dalgarno or translational start site of an mRNA, precluding the ribosome and in some cases resulting in destabilization of the mRNA. To determine the potential for PrrH to directly regulate expression of proteins involved in pyochelin biosynthesis or uptake, we analyzed the H-IG sequence to determine if pairing could occur with any of the PrrH affected *pch* mRNAs using CopraRNA (54-56). This analysis identified PrrH complementarity sites within the coding sequences of two distinct pyochelin genes: *pchI* and *pchE* (**Fig. 5A**), demonstrating the capacity of the PrrH-IG sequence to directly pair with at least two mRNAs encoding proteins for pyochelin biosynthesis. Of note, the *pchI* and *pchE* are located within a single operon (**Fig. 5A**), suggesting PrrH may be able to bind at two distinct sites of the *pchEFGHI* mRNA.

**Fig 5.**
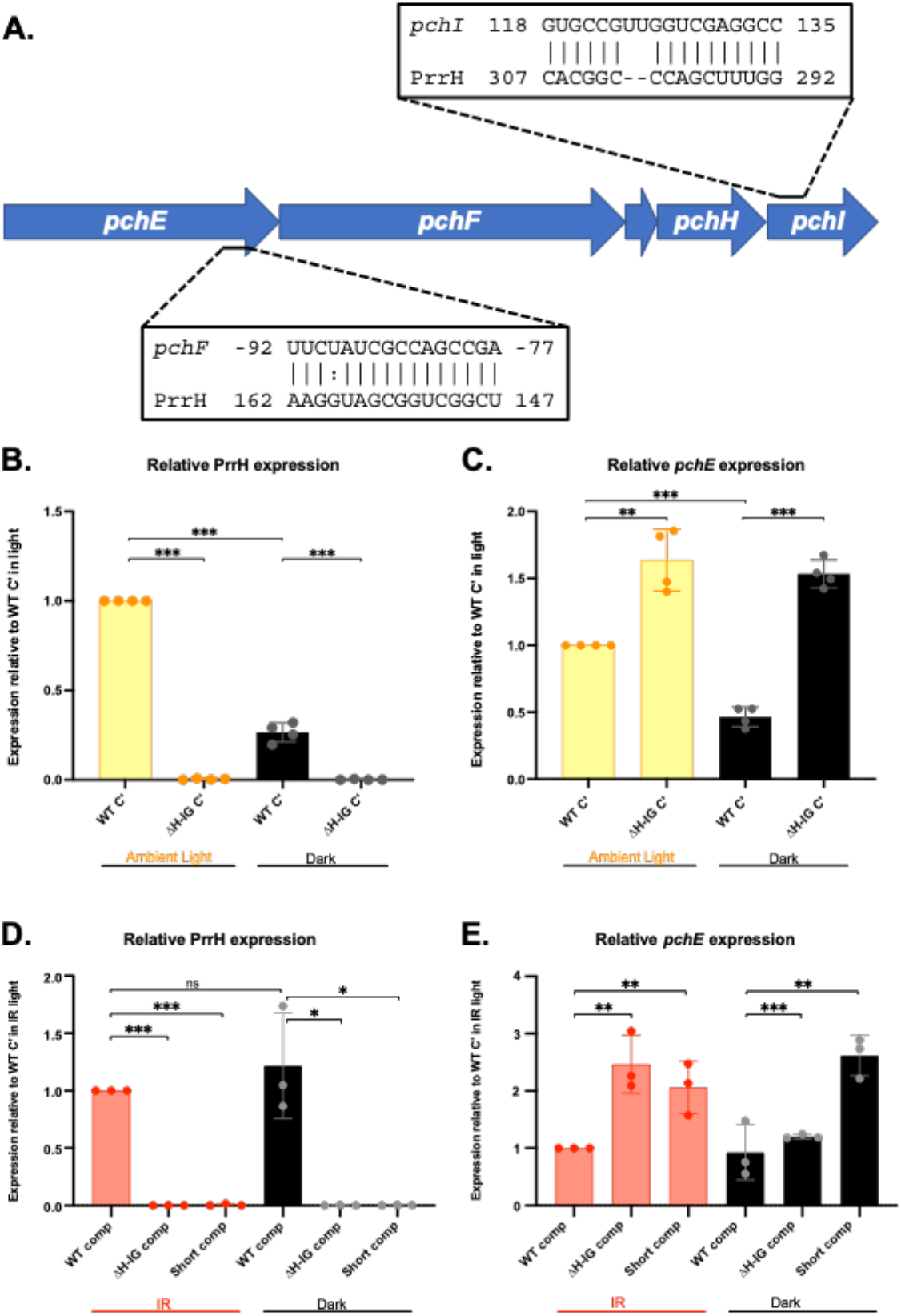
Static growth and controlling for light promote consistent PrrH repression of the *pchEF* mRNA. (A) CopraRNA identified sequences with complementarity to PrrH upstream of *pchF* and within *pchI*. (B-E) qRT-PCR analysis of the PrrH sRNA (B,D) and *pchE* mRNA (C,E) levels relative to WT-comp shown as an average of 3 or 4 biological replicates. RNA was isolated from cultures grown in M9 media supplemented with 50 nM FeCl_3,_ grown in static conditions at 37°C for 8 hours, and exposed to either ambient or infrared (IR) light as described in the materials and methods. Expression is calculated as relative to WT-comp in static, light conditions. Significance was calculated using a two-tailed Student’s t test with asterisks indicating the following P values: * indicates P<0.05, ** P<0.005, and *** P<0.0005.

We next sought to determine whether RNA levels for pyochelin biosynthesis are affected by PrrH mutation under conditions where this system is more strongly expressed. Recently, our lab observed that proteins involved in pyochelin biosynthesis are more robustly regulated by iron in static compared to shaking conditions (57), suggesting static conditions may be more permissive for expression of pyochelin mRNAs. Therefore, we performed qRT-PCR on samples collected from both static and shaking cultures and, indeed, we observed an increase in *pchE* transcription in each *prrH* mutant compared to the WT-comp when grown in static, but not shaking, conditions (**Supplementary Materials Fig. S4A**). However, the increase in *pchE* expression, albeit statistically significant, was modest, and results of this experiment when repeated several weeks apart gave inconsistent results (**Supplementary Materials Fig. S4B**). Thus, while these data suggested that PrrH may affect *pchE* mRNA stability, they also indicated that confounding variables were continuing to affect our ability to study PrrH regulation.

### Ambient and infrared light affect PrrH’s impact on *pchE* mRNA levels

Recent studies from Mukherjee, *et al*, demonstrated that the photoreceptor BphP can mediate light-dependent changes in biofilm-related gene expression via the response regulator AlgB (58). The BphP photoreceptor activity is dependent on a biliverdin chromophore produced via the BphO heme oxygenase, which serves independent functions from the HemO heme oxygenase that mediates acquisition of iron from heme (59, 60). Owing to its potential role in heme homeostasis, we wondered whether BphP, and therefore light, may affect PrrH regulation of the potential *pchE* target. To begin testing this, we first assessed the impact of ambient light on static cultures of the WT-comp and ΔH-IG comp strains. Cultures were grown in a well-lit room (fluorescent lights on and near a window) to provide an “ambient light” source, and parallel cultures were wrapped completely in foil to produce a “dark” condition. Initial qPCR analysis revealed a significant decrease in PrrH expression in dark compared to light conditions (**Fig. 5B**), though this trend was reversed in a subsequent experiment (**Supplementary Materials Fig. S4C**). In contrast, we noted a significant increase in *pchE* expression upon H-IG deletion in both light and dark conditions (**Fig. 5C**), a finding that was reproduced in a subsequent run of the experiment (**Supplementary Materials Fig. S4D**).

We next determined the impact of infrared (IR) light, which is specifically detected by the BphP phytochrome, on static cultures of the WT comp, ΔH-IG comp, and short-comp strains. IR light had no impact on expression of PrrH levels (**Fig. 5D**). However, IR light allowed for robust induction of *pchE* expression upon *prrH* mutation (**Fig. 5E**). While the mechanistic rationale for the effects of light on PrrH and *pchE* expression remain poorly understood, these experiments demonstrate how controlling for light as an environmental variable may be critical to studying the function of this unique sRNA.

### Heme negatively affects *pchE* expression

Previous work demonstrates that heme positively affects PrrH sRNA levels via the PhuS heme binding protein (50). We therefore determined if heme also affected expression of *pchE* in the WT PAO1 strain, lacking the pucP18 vector, in static conditions either in the presence of IR light or wrapped in foil. At three hours post-inoculation, one of each duplicate culture was supplemented with 5 µM heme, and cultures were grown for another 8 hours. Heme supplementation had a positive impact on PrrH levels, both in dark and IR light conditions (**Fig. 6A**); however due to variability in PrrH expression, the increase was only statistically significant when cultures were incubated in the dark (**Fig. 6A**). Notably, we observed a robust and statistically significant decrease in *pchE* levels upon heme supplementation, both in IR and dark conditions (**Fig. 6B**). We did not observe any repression of the PrrF sRNAs (**Fig. 6C**), suggesting that heme repression of *pchE* is not due to Fur-mediated repression from the iron that is enzymatically released upon heme degradation. Thus, our data suggest that heme specifically represses expression of the *pchE* gene in WT PAO1.

**Figure 6.**
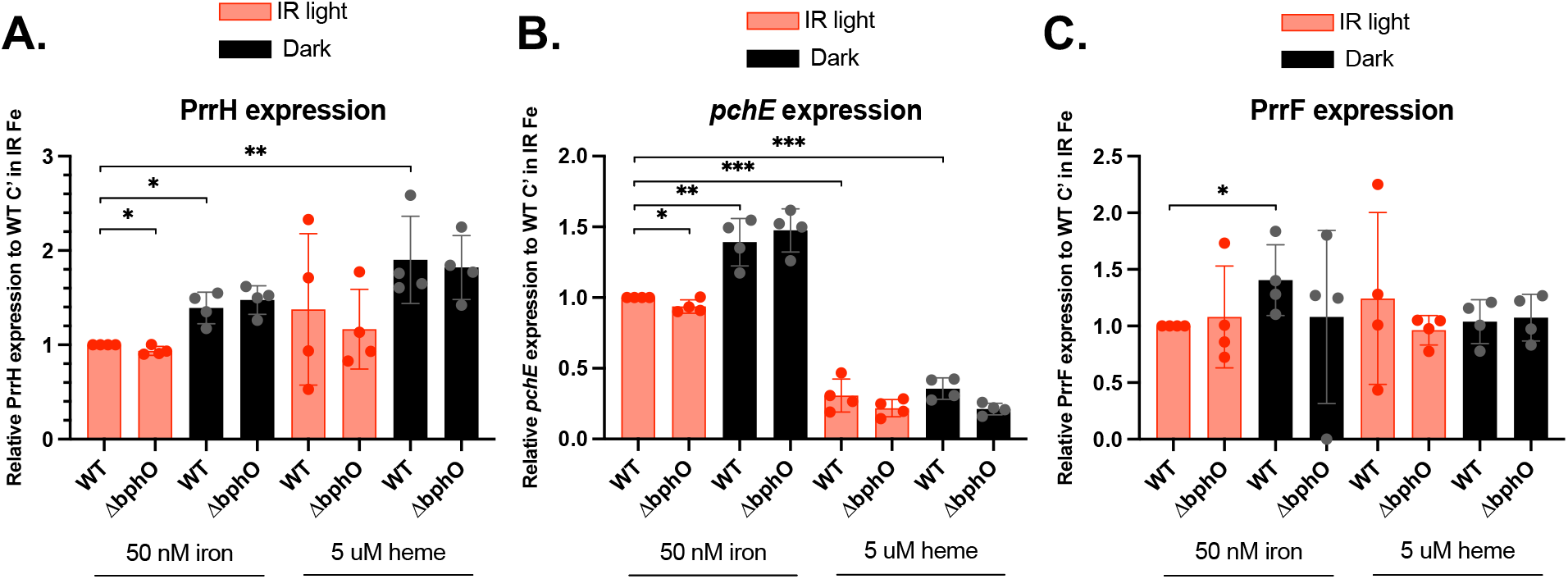
Heme negatively affects *pchE* expression. Relative PrrH (A), *pchE* (B), and PrrF (C) transcript levels relative to WT PAO1 when grown in the presence of absence of infrared (IR) light. Data are an average of 4 biological replicates in M9 media supplemented with 50 nM FeCl_3_ or 5 µM heme grown statically at 37°C for 8 hours. Significance was calculated using an unpaired *t* test with two-tailed P values where * indicates P<0.05, ** P<0.005, and *** P<0.0005.

We next determined if heme repression of *pchE* was dependent on the PrrH sRNAs using the WT-comp, ΔH-IG, and short *prrF* complementation strains. The three strains were grown in static conditions, with or without heme supplementation as above, and analyzed for PrrH, *pchE*, and PrrF expression. We noted a similar positive but statistically insignificant effect of heme supplementation on PrrH expression in this experiment (**Supplementary Materials, Fig. S5A**). Surprisingly, we did not see any repression of *pchE* by heme in the WT comp strain, though heme did appear to eliminate the induction of *pchE* in the *prrH* mutants (**Supplementary Materials, Fig. S5B**). It is not clear why the heme regulation of *pchE* is not evident in the WT comp strain, though we did note in our omics studies (above) that iron regulatory pathways are somewhat altered in the complemented strains. Thus, we cannot assert, at this time, the role of PrrH in heme-mediated repression of *pchE*. However, our data do indicate a role for both heme and PrrH in affecting expression of *pchE*.

### BphO is not required for PrrH- or heme-mediated regulation of *pchE*

Since our data suggest that IR light may affect reproducibility of PrrH regulation on *pchE*, we next sought to determine if the Bph phytochrome system affected expression of either PrrH or *pchE*. For this we used a deletion mutant of the gene encoding the BphO heme oxygenase, which generates α-biliverdin (α-BVIX) as the chromophore for the BphP phytochrome. The Δ*bphO* mutant showed similar expression of PrrH, *pchE*, and PrrF under all conditions tested, regardless of IR light exposure (**Fig. 6**). Thus, the impact of light on eliminating variability of this regulatory pathway does not seem to be directly due to the BphP phytochrome. Instead, light may affect other metabolic pathways that indirectly affect the ability of heme to affect expression of PrrH and *pchE*.

## Discussion

This study aimed to determine the regulatory impact of the heme-responsive PrrH sRNA in *P. aeruginosa*. Heme has emerged as a significant factor in chronic *P. aeruginosa* infections over the past decade, with multiple studies demonstrating a reduced reliance on siderophore mediated iron uptake in the CF lung (33-37). Yet, only a few studies have rigorously addressed the global impacts of heme on *P. aeruginosa* gene expression. Here, we characterized two distinct *prrH* mutants, which express wild type levels of the PrrF sRNAs but produce truncated PrrH sRNAs, and we determined the impact of these mutants on global gene expression using RNASeq and proteomics. While experimental variations were observed in PrrH regulons from separate experiments, we were able to identify pyochelin as a consistently dysregulated gene in the *prrH* mutants and therefore a potential target of the PrrH sRNA. We further determined that static growth promoted PrrH-dependent changes in *pchE* RNA levels, and we identified ambient light as a likely confounding variable in our early studies. The impact of light builds on the recent discovery that BphP, an α-biliverdin dependent photoreceptor (59), controls activity of the AlgB response regulator in *P. aeruginosa (58)*. This discovery led us to consider how light might affect

PrrH-dependent heme regulation, allowing us to control for this variable in the current study. The result is the identification of *pchE* as the first verified PrrH-responsive gene in *P. aeruginosa*.

To our knowledge, a link between heme regulation and siderophore synthesis has only been shown in one other bacterial species, *Staphylococcus aureus* (61). The *S. aureus* staphyloferrin B (Sbn) biosynthetic locus contains a gene encoding the SbnI protein, which induces expression of the *sbn* operon in its apo form, and functions in Sbn synthesis when bound to heme (61). Thus, SbnI plays a bifunctional role in siderophore biosynthesis and modulating heme-dependent regulation of siderophore gene expression. The *P. aeruginosa* PhuS protein similarly appears to have a bifunctional role, as it binds to the *prrH/prrF1* promoter in its apo form, and functions as a shuttle to HemO when bound to heme (50, 60, 62, 63). Here, we show that PrrH negatively impacts expression of *pchE*, and we show that heme similarly blocks *pchE* gene expression in *P. aeruginosa*. These results highlight heme as a preferred iron source under static growth conditions, which promotes biofilm intiation and may more closely mimic chronic biofilm infections where heme is a preferred iron source.

The impact of heme uptake and regulation on *P. aeruginosa* virulence has become increasingly appreciated in the past decade. Recent work demonstrated a role for the unique β and δ biliverdin isomers in post-transcriptional regulation of the Has heme uptake system (64), which is required for *P. aeruginosa* virulence (51). Expression of the Has system is also subject to heme binding to the secreted HasA hemophore, which binds to the HasR outer membrane receptor. This, in turn, initiates a cell surface signaling (CSS) cascade through the HasI extracytoplasmic function (ECF) sigma factor and cognate HasS anti-sigma factor (64). As indicated above, PhuS functions as another heme-dependent regulatory protein, by binding in its apo form to the *prrF1/prrH* promoter to affect PrrH expression (50). Heme has additionally been implicated in virulence gene regulation via the BphP photoreceptor, which requires α-biliverdin produced by the BphO heme oxygenase (58, 59). This recent study showed that AlgB acts as the response regulatory to the BphP sensor kinase activity when sensing light, allowing light to negatively impact biofilm formation (58). Notably, heme degradation by BphO does not contribute to *P. aeruginosa’s* ability to use heme as an iron source, but instead seems to turn over intracellularly produced heme (60, 62). It is, therefore, intriguing that light was a confounding environmental variable in our studies of PrrH regulation, suggesting that these heme-dependent regulatory systems are interlinked. Our studies suggest that the Bph system does not directly impact expression of PrrH or *pchE* (**Fig. 6)**. Studies of IR light regulation via BphP revealed a large metabolic regulon (58), which may in turn indirectly influence many aspects of *P. aeruginosa* iron and heme homeostasis. Studies into how the PrrH heme regulatory system intersect with photosensing are continuing in our groups.

Understanding how heme regulation functions in *P. aeruginosa* has been complicated by additional factors, including the transient presence of heme as an extracellular iron source. Once extracellular heme is transported into the cell, it is shuttled to HemO to be degraded to biliverdin which, as described above, exerts its own regulatory effects on gene expression. Heme oxygenase also releases iron that can then function through the Fur protein to repress expression of heme-responsive genes, including *hasR* and *prrF1/prrH*. Thus, time-dependent observations of heme flux, promoter activity, RNA levels, and protein expression are likely all critical for understanding heme-dependent regulatory effects. Notably, supplementation of cultures with higher concentrations of heme (≥ 20 µM) has been pursued in other works, yet these levels have the potential to initiate toxicity toward cell membranes, nonspecific oxidative-cleavage prior to uptake, and the formation of μ-oxo-dimers resulting in stable and biologically unavailable heme polymers. In the current study, we began assessing the impact of heme on PrrH and *pchE* levels during static growth, while all previous heme regulatory studies have been conducted in shaking growth. Similar to observations by our group regarding the impacts of static growth on global iron regulation (47), our work here suggests that heme and PrrH regulation is altered under static conditions. Static culture conditions result in slower growth of *P. aeruginosa*, which will require us to reassess the timing of heme uptake and metabolism to release iron under these conditions. Static cultures also likely result in heterogenous communities with varying heme uptake and regulatory activities. Continued work on heme uptake and regulation in complex *P. aeruginosa* communities, such as within static cultures and biofilm communities, is required to understand more physiologically relevant implications of these regulatory pathways.

An additional complexity for studies described here is the overlapping sequence of the PrrF and PrrH sRNAs. Two recent studies ascribed a variety of physiological functions to PrrH, yet both studies used knockouts and complementation constructs containing the entire *prrF-prrH* sequence (65, 66). Therefore, any observations or phenotypes derived from these strains cannot simply be designated as specific to PrrH. The genetic strategy previously developed by Reinhart, *et al* (67) allowed us to overcome this issue by focusing on the sequence that is unique to PrrH. We note that this system is not ideal because PrrF and PrrH are overexpressed in these strains, leading to some changes in global gene expression (**Fig. S1, 3**), and indeed the WT comp showed a different effect of heme on expression of *pchE* (**Fig. S4, 6**). We continue to work toward genetic strategies that will allow us to specifically affect PrrH function while allowing the PrrF sRNAs to remain unaffected.

Overall, this study identified the first regulatory target of the PrrH sRNA as *pchE* and provided evidence that this regulation could occur through direct post-transcriptional regulation of the *pchE* mRNA. Additionally, we provide evidence that light may affect systems involved in heme regulation, necessitating careful control of this environmental condition for our studies. Current and future work will examine time-dependent effects of heme supplementation on these systems, and work toward identifying more global impacts of the PhuS and PrrH regulatory molecules.

## Materials & Methods

### Bacterial strains and growth conditions

Strains used in this study are listed in **Table S1**. *P. aeruginosa* strains were maintained on LB or BHI agar or broth. Strains carrying the complementation plasmids were maintained with carbenicillin (250 µg/mL). Media was supplemented with iron or heme as follows: 50 nM FeCl_3_ (−Fe), 100 µM FeCl_3_ (+Fe), and 1 µM or 5 µM Heme (+He) prepared as previously described (44). Overnight cultures were grown in LB or BHI broth, aerobically (250 RPM, 37°C) and washed in M9 media (Teknova, Hollister, CA) prior to inoculating into M9 supplemented with iron or heme to a starting OD_600_=0.05. Shaking cultures were grown in 1:10 (media:flask) ratios, shaking aerobically at 250 RPM, 37°C and collected at 8 hours for analyses. Static cultures were grown in 24-well cell culture plates (Greiner Bio-One, Kremsmunster, Austria) at 37°C and collected at 8 hours for analyses. For ambient light conditions, cultures were grown in a well-lit room (fluorescent overhead lights and near a window during the day). For infrared (IR) light conditions, plates were placed under a 730 nM LED Lightbar (Forever Green Indoors, Inc, Seattle, WA). Plates were wrapped in foil for dark conditions. For RNA isolation, cultures were mixed with an equal volume of RNALater (Sigma-Aldrich, St. Louis, MO) and stored at -80°C until processing.

### Quantitative real-time PCR (qRT-PCR)

RNA was isolated following manufacturer’s suggested protocol using the RNeasy Mini Kit (Qiagen, Hilden, Germany). An additional DNase I (New England Biolabs, Ipswich, MA) treatment was performed at 37°C for 2 hours, ethanol precipitated, and eluted in RNase-free water. qRT-PCR was performed as previously described (43, 44). Primer and probe sequences are listed in **Table S2**.

### Northern blot analyses

RNA was isolated following manufacturer’s suggested protocol using the RNeasy Mini Kit (Qiagen, Hilden, Germany). 5 µg (PrrF1, PrrF2) or 20 µg (PrrH) of total RNA was electrophoresed on a 10% denaturing urea TBE gel (Bio-Rad, Hercules, CA). The RNA was then transferred to a BrightStar-Plus Positively Charged Nylon Membrane (Invitrogen, Carlsbad, CA) using a Trans-Blot Turbo Transfer System (Bio-Rad) and crosslinked for 2 minutes using a UV Crosslinker (VWR, Radnor, PA). The blots were incubated with ^γ32^-P 5’-labelled probe at 42°C overnight and imaged using a phosphor screen on a Typhoon FLA 7000 Variable Mode Imager System (GE Healthcare, Chicago, IL). Probe sequences are listed in **Table S2**.

### RNAseq

Cultures were grown in shaking conditions as described above, without any additional controls for light. Cultures were collected (at 8 hours of growth) directly into RNALater. Iron was supplemented at a concentration of 50 nM (low iron) or 100 µM (high iron), while heme was used at 5 µM. Sample preparation and subsequent RNA extraction and analyses were performed as previously described (51). RNA integrity was validated using an Agilent 2100 Bioanalyzer. Libraries were prepared with samples with an RNA Integrity Number (RIN) greater or equal to 8. Ribosomal RNA was depleted using the Ribo Zero kit and samples were converted into Illumina sequencing libraries using the ScriptSeq v2 RNA-Seq Library Preparation Kit (Epicentre, Illumina). Libraries were sequenced using Illumina HiSeq (2 × 150 bp reads). Three biological replicates were sequenced in each group and an average of 40 million reads were obtained for each sample. Reads were mapped against the reference genome of *P. aeruginosa* PAO1 (NC_002516) with the following settings: mismatch cost = 2, insertion cost = 3, deletion cost = 3, length fraction = 0.8, similarity fraction = 0.8. Fold changes in gene expression, and statistical analyses were performed using the extraction of differential gene expression (EDGE) test as implemented in CLC Genomics, which is based on the Exact Test. Differential gene expression was calculated by comparing samples and setting a fold-change cut-off as described in the figure legends. Genes were included in further analysis only if differences in expression yielded a FDR *p-*value of *p* ≤ 0.05.

### Quantitative label-free proteomics

Cultures were grown in shaking conditions as described above, without any additional controls for light. Cultures were collected at 8 hours of growth. Iron was supplemented at a concentration of 50 nM (low iron) or 100 µM (high iron), while heme was used at 5 µM. Sample preparation and subsequent proteomics analyses were performed as previously described (47, 68, 69). In brief, cells were harvested by centrifugation at 2000 rpm for 30 s at 4 °C, and then subsequently lysed in 4% sodium deoxycholate, reduced, alkylated, and trypsinolyzed on filter as previously described (70). Tryptic peptides were separated on a nanoACQUITY UPLC analytical column (BEH130 C_18_, 1.7 μm, 75 μm x 200 mm, Waters) over a 165 min linear acetonitrile gradient (3-40%) with 0.1 % formic acid using a Waters nanoACQUITY UPLC system and analyzed on a coupled Thermo Scientific Orbitrap Fusion Lumos Tribrid mass spectrometer. Full scans were acquired at a resolution of 120,000, and precursors were selected for fragmentation by higher-energy collisional dissociation (normalized collision energy at 30%) for a maximum 3-second cycle. Tandem mass spectra were searched against the *Pseudomonas* genome database PAO1 reference protein sequences (71) using the Sequest-HT and MS Amanda algorithms with a maximum precursor mass error tolerance of 10 ppm (72, 73). Carbamidomethylation of cysteine and deamidation of asparagine and glutamine were treated as static and dynamic modifications, respectively. Resulting hits were validated at a maximum false-discovery rate (FDR) of 0.01 using the semi-supervised machine learning algorithm Percolator (74). Protein abundance ratios were measured by comparing the MS1 peak volumes of peptide ions, whose identities were confirmed by MS2 sequencing. Label-free quantification was performed using an aligned AMRT (Accurate Mass and Retention Time) cluster quantification algorithm (Minora; Thermo Fisher Scientific, 2017). Protein interactions were analyzed using STRING 10.5 and visualized with Cytoscape 3.8.0 (75, 76)

## Acknowledgements

We thank Dr. Susana Mouriño for her assistance in establishing appropriate conditions for heme regulation in these studies. We also thank Prof. Beronda Montgomery for insightful conversations about the impact of light in photoreceptors in non-photosynthetic bacteria. This work was funded by NIH grants R01-AI23320 (AGO), AI161294 (AW and AGO), and the University of Maryland School of Pharmacy Mass Spectrometry Center (MAK).

